# FedEdgeR: federated and privacy-preserving edgeR for differential gene expression analysis

**DOI:** 10.64898/2026.09.17.752480

**Authors:** Xun Song, Zhi Wei

## Abstract

**Motivation:** Multi-center RNA-seq studies improve statistical power, but privacy regulations restrict patient-level data sharing. Meta-analysis methods avoid this restriction but lose power, especially under per-site imbalance. Federated learning allows sites to share only summary statistics. Among the three dominant differential expression (DE) frameworks, Flimma federates limma-voom and FedPyDESeq2 federates DESeq2. edgeR, still preferred for small-sample and high-variability designs, lacks a federated counterpart. Its iteratively reweighted least squares (IRLS) and Cox-Reid dispersion estimation make federation harder than limma-voom’s single-pass fit.

**Results:** We present FedEdgeR, a federated edgeR implementation protected by secure multi-party computation (SMPC), covering IRLS GLM fitting, three-level dispersion estimation, and the likelihood-ratio test. We evaluate it on four representative RNA-seq datasets: two tumor/normal cohorts, a multi-cohort anti-PD-1 immunotherapy study, and a 6-sample paired stress-test. FedEdgeR matches pooled edgeR under various metrics, such as Pearson *r* ≥ 0.99999 on − log_10_ *p*-values, 100% complete top-100 DE gene overlap, and *F*_1_ = 1.000 at the nominal FDR level 0.05.

It consistently outperforms Fisher, Stouffer, random-effects, and RankProd meta-analysis on every tested dataset.

**Availability:** Source code is archived at https://doi.org/10.5281/zenodo.20450301.

## 1 Introduction

Differential expression (DE) analysis is fundamental to characterizing transcriptomic responses to pathological, clinical, and environmental factors (Wang et al., 2009). Beyond RNA-seq, the same statistical approaches have been applied to other comparative high-throughput sequencing (HTS) assays (Love et al., 2014). Three statistical frameworks currently dominate the field: edgeR, DESeq2, and limma-voom. EdgeR fits negative binomial generalized linear models (GLMs), and employs empirical Bayes shrinkage to estimate gene-level dispersions (Robinson et al., 2009; McCarthy et al., 2012). DESeq2 adopts a similar GLM framework but uses a different shrinkage strategy for dispersion estimation (Love et al., 2014). In contrast, limma-voom transforms count data to log-CPM values and applies weighted linear models (Ritchie et al., 2015; Law et al., 2014). Of these three, edgeR was the first specifically designed for sequencing count data (Robinson and Smyth, 2007). It remains a preferred method for datasets with small sample sizes or high biological variability (Robinson and Smyth, 2007; Chen et al., 2025).

Combining samples from multiple medical centers or research sites can increase the statistical power of DE analyses. Larger and more diverse cohorts help reduce batch-specific biases and improve the generalizability of results (Leek et al., 2010). However, sharing patient-level gene expression data across institutions is difficult in practice. Privacy regulations, such as the General Data Protection Regulation (GDPR), impose strict constraints on the transfer of identifiable genomic data (Brauneck et al., 2024). Even summary statistics, such as allele frequencies, can be used to re-identify individuals under certain conditions (Homer et al., 2008). These legal and technical limitations create a tension between the pursuit of large-scale, multi-site genomic studies and the need to protect patient privacy.

The most common workaround is meta-analysis. Each site runs a DE analysis on its own data. The per-site p-values or effect sizes are then combined using methods such as Fisher’s method (Fisher, 1934), Stouffer’s method (Stouffer et al., 1949), or the DerSimonian-Laird random-effects model (RE model) (DerSimonian and Laird, 1986). This approach avoids raw data sharing, but treats each site as an independent study and therefore loses statistical power relative to a pooled analysis. The loss is more severe when sample sizes differ across sites or when one site has a biased ratio of cases to controls (Zolotareva et al., 2021; Nasirigerdeh et al., 2022). When per-site sample size approaches the design rank, the GLM is saturated and the per-site dispersion is no longer identifiable. No combiner can recover this lost information.

Federated learning provides an alternative. In a federated analysis, each site keeps its raw data locally and only shares intermediate summary statistics with a central server (McMahan et al., 2017; Kairouz and McMahan, 2021). The server combines these statistics to produce a global result that is mathematically equivalent to a pooled analysis. When combined with secure multi-party computation (SMPC), even the intermediate statistics can be protected from the server (Bonawitz et al., 2017). Several federated frameworks for biomedical data now exist, including HyFed (Nasirigerdeh et al., 2021), FeatureCloud (Matschinske et al., 2023), and DataSHIELD (Gaye et al., 2014).

Two federated tools for RNA-seq DE analysis have been proposed. Flimma (Zolotareva et al., 2021) federates the limma-voom pipeline using HyFed’s three-party SMPC architecture. Because limma-voom fits each gene by weighted least squares (WLS), Flimma requires only one round of secure aggregation per gene. FedPyDESeq2 (Muzellec et al., 2024) federates the DESeq2 pipeline, but it does not include cryptographic privacy protection. Although edgeR is still widely used in small-sample and high-variability RNA-seq DE analysis, no federated version exists. Federating edgeR poses challenges that do not arise when federating limma-voom or DESeq2. First, edgeR fits negative binomial GLMs through iteratively reweighted least squares (IRLS). This requires multiple rounds of secure communication per gene, rather than the single pass used by limma-voom. Second, edgeR estimates dispersions at three levels using the Cox-Reid adjusted profile likelihood (APL) (Cox and Reid, 1987). Each evaluation of this likelihood calls the IRLS procedure internally, which raises both the communication and the privacy cost.

We present FedEdgeR, the first federated implementation of edgeR’s DE pipeline. The implementation covers the IRLS GLM fit, the three-level dispersion estimation, and the likelihood-ratio test. FedEdgeR makes three contributions:

- It is the first federated DE pipeline to combine SMPC (Bonawitz et al., 2017) privacy protection with nonlinear iterative model fitting. Flimma’s SMPC protects only a single-pass linear model, and FedPyDESeq2 federates DESeq2 without cryptographic protection.
- We close the federation gap for edgeR’s DE pipeline: the IRLS GLM fit, three-level dispersion estimation, and Cox-Reid APL. A complete mathematical derivation shows that each IRLS iteration’s sufficient statistics (**X**^*T*^ **WX** and **X**^*T*^ **Wz**) are additive across sites and can be securely aggregated.
- On four RNA-seq datasets, the Pearson correlation of *p*-values with pooled edgeR is ≥ 0.99999 and the F1 score at FDR *<* 0.05 is 1.000. The common, trended, and tagwise dispersions all match the centralized pipeline. FedEdgeR outperforms four meta-analysis methods on all datasets.

## 2. Methods

### 2.1 System architecture and privacy model

FedEdgeR uses the HyFed hybrid federated architecture (Nasirigerdeh et al., 2021), which involves three parties: *K* client sites, one aggregator, and one compensator. Each client holds a local count matrix **Y**^*i*^ and a design matrix **X**^*i*^ for *i* = 1, …, *K*. The aggregator coordinates the computation and stores no raw data. The compensator is a separate server that sums the random noise terms from each client.

When a client needs to send a local statistic **M**^*i*^ to the aggregator, it first generates a random noise matrix **N**^*i*^ of the same shape. The client sends the masked value **M**^*i*^ + **N**^*i*^ to the aggregator and the noise **N**^*i*^ to the compensator. The compensator sums all noise terms into a global noise 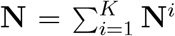 and sends it to the aggregator. The aggregator then recovers the true global sum:

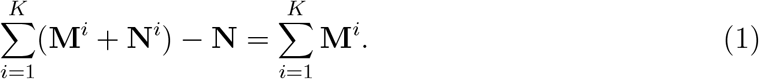

For integer parameters, the noise uses uniform sampling over a finite field ℤ_*p*_ with modular arithmetic. For real-valued parameters, the noise is Gaussian with *σ*^2^ = 10^12^ (Nasirigerdeh et al., 2021) (Supplementary Note S1). Following the threat model of Flimma and sPLINK (Zolotareva et al., 2021; Nasirigerdeh et al., 2022), all parties (clients, aggregator, compensator) are honest-but-curious and do not collude.

### 2.2 Federated edgeR algorithm

FedEdgeR’s federation contribution covers edgeR’s DE pipeline: the IRLS GLM fit, the three-level dispersion estimation, and the likelihood-ratio test. The FedEdgeR pipeline follows the same sequence of steps as centralized edgeR (McCarthy et al., 2012; Chen et al., 2025). Figure 1 gives an overview of the pipeline. At each step, clients compute local summary statistics from their own data and send them to the aggregator through SMPC masking. The aggregator combines these statistics into updated parameters. Within an iterative step, it broadcasts the new parameters back to the clients for another round; once the step has converged, the pipeline moves on.

**Figure 1:**
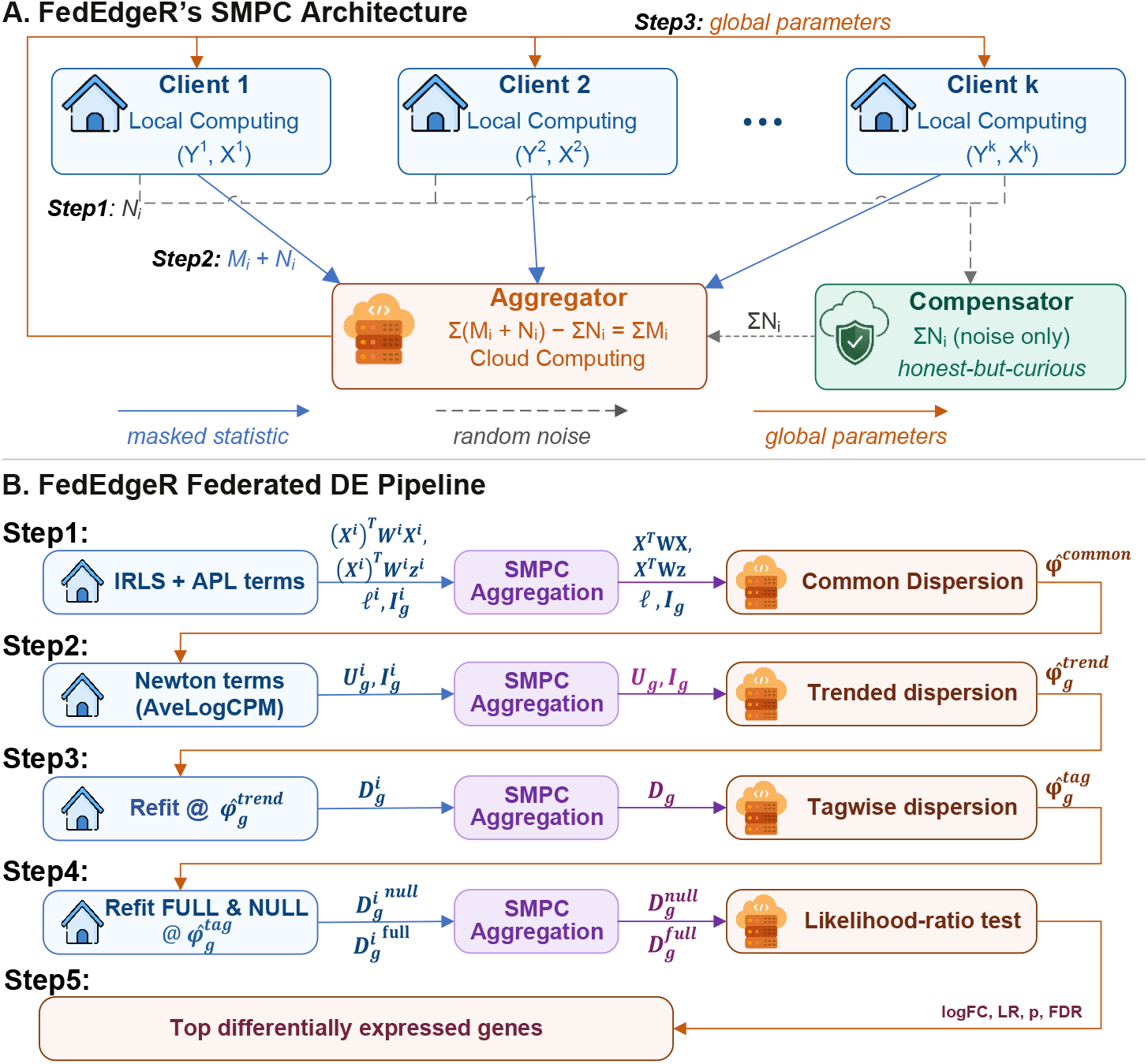
FedEdgeR architecture and DE pipeline. **(a)** SMPC mask-and-denoise architecture. Each client sends a masked statistic *M*_*i*_ + *N*_*i*_ to the aggregator and the raw noise *N*_*i*_ to the compensator. The aggregator recovers ∑_*i*_ *M*_*i*_ without ever seeing any individual *M*_*i*_. Raw counts and design matrices never leave the client sites. **(b)** FedEdgeR federated DE pipeline. Each row is split by color into client local computation (blue), SMPC aggregation (green), and aggregator inference (orange). Each box lists only the quantities computed and transmitted at that stage, not the full party state. Broadcast parameters (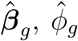, dispersion estimates) and earlier SMPC-aggregated outputs persist at the receiver between stages and are reused without retransmission. **Alt text:** Two-panel diagram. Panel (a): three-party SMPC architecture with *K* client sites, one aggregator, and one compensator. Clients send masked statistics *M* + *N* to the aggregator and the raw noise *N* to the compensator, and the aggregator subtracts the compensator’s noise sum to recover the global sum of *M* without seeing individual client values. Panel (b): federated DE pipeline arranged as five rows, each row split by color into client local computation (blue), SMPC aggregation (green), and aggregator inference (orange). The rows output common, trended, and tagwise dispersions, the likelihood-ratio test, and the BH-FDR-ranked DE gene list.

#### 2.2.1 Normalization and filtering

The pipeline takes per-sample offsets as input parameters and is agnostic to how those off-sets are produced. Each offset is the log of an effective library size, defined as the product of raw library size and normalization factor. Our benchmarks use edgeR’s default TMM normalization (Robinson and Oshlack, 2010) computed centrally on the merged per-site count matrices. This lets pooled edgeR and FedEdgeR share the same upstream input, so the equivalence claim isolates the federated DE pipeline. Deployments requiring end-to-end privacy can swap in Flimma’s federated upper-quartile normalization (Zolotareva et al., 2021; Bullard et al., 2010) without any change to the downstream pipeline. The `filterByExpr` step likewise has a federated form in Flimma.

#### 2.2.2 GLM fitting with federated IRLS

EdgeR fits a negative binomial (NB) generalized linear model (GLM) to each gene *g* via iteratively reweighted least squares (IRLS) (McCarthy et al., 2012). Under this model, the count *y*_*gs*_ for gene *g* in sample *s* is modeled as *y*_*gs*_ ~ NB(*µ*_*gs*_, *ϕ*_*g*_), with mean 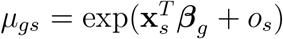 and variance Var(*y*_*gs*_) = *µ*_*gs*_(1 + *µ*_*gs*_*ϕ*_*g*_). Here **x**_*s*_ is the row of the design matrix for the sample *s, o*_*s*_ is the offset of the sample (Section 2.2.1), and *ϕ*_*g*_ is the dispersion. At each IRLS iteration *t*, the algorithm computes a diagonal weight matrix **W**_*g*_ and a working variable **z**_*g*_ from the current ***β***_*g*_ :

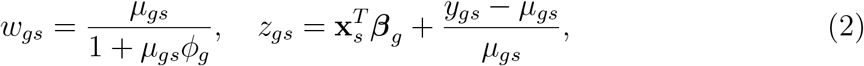

where the offset enters only through *µ*_*gs*_ (Eq. 2 keeps *z*_*gs*_ free of an explicit offset term). The weighted least-squares update below therefore returns ***β***_*g*_ directly without an offset subtraction. The coefficient vector is then updated by:

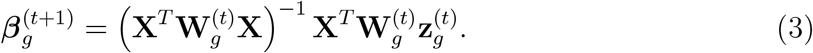

Both **X**^*T*^ **W**_*g*_**X** and **X**^*T*^ **W**_*g*_**z**_*g*_ can be expressed as additive decompositions over individual samples. Since the datasets are distributed among *K* clients, we can reformulate these terms as follows:

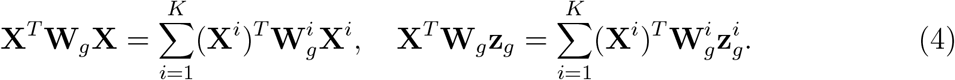

Each client computes its local terms using only its own data and the current ***β***_*g*_ (broadcast by the aggregator). These local terms are then transmitted to the aggregator using the SMPC masking described in Section 2.3. The aggregator sums them, solves for the new ***β***_*g*_, and broadcasts it back. This process repeats until ***β***_*g*_ converges. We process all genes in parallel within each iteration to reduce the number of SMPC rounds.

#### 2.2.3 Dispersion estimation

Consistent with edgeR, FedEdgeR supports multiple dispersion estimation strategies and allows for common, trended, and tagwise (gene-specific) dispersion values across the distributed datasets (McCarthy et al., 2012; Robinson and Smyth, 2007). The three estimators are detailed below according to their computational dependency. The full federated procedure is given in Supplementary Note S6.

##### Common dispersion

A straightforward approach is to assume a common dispersion across all genes, such that *ϕ*_*g*_ = *ϕ* for all *g* (McCarthy et al., 2012; *Robinson and Smyth, 2007*). *FedEdgeR searches over a grid of 21 candidate values spaced evenly on the log scale. For each candidate ϕ*, we run the federated IRLS to fit the GLM and then evaluate the Cox-Reid adjusted profile likelihood (APL) (Cox and Reid, 1987). Each client computes its local negative-binomial log-likelihood 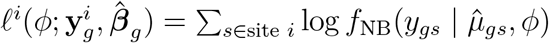 and Fisher information 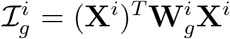. These are aggregated via SMPC, and the aggregator evaluates

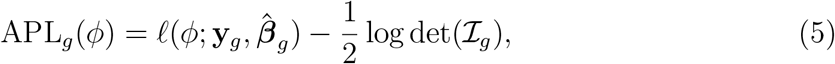

where 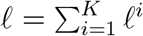 is the global NB log-likelihood, 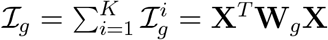 is the Fisher information matrix, and the 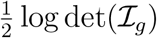 term is the Cox-Reid penalty that corrects the bias from profiling out ***β***_*g*_. Throughout the APL framework, *ϕ* is the dispersion at which the profile likelihood is evaluated, and 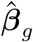 is the maximum-likelihood estimate of the coefficients at that *ϕ*. The common dispersion is then 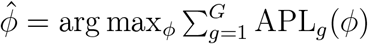. The maximum is located by a natural cubic-spline interpolant through the 21 grid points (see Supplementary Notes S4 and S5 for the federated GLM and APL derivations).

##### Trended dispersion

Beyond common dispersion, *ϕ*_*g*_ may vary smoothly with the average expression of each gene. We first compute each gene’s AveLogCPM through a federated Fisher-scoring iteration on the NB intercept-only mean model (McCarthy et al., 2012; Chen et al., 2025) with a fixed plug-in *ϕ*_*g*_ = 0.05, the edgeR aveLogCPM default. We then apply local-regression smoothing on the (AveLogCPM, log *ϕ*_*g*_) plane using R’s locfit (Loader, 1999), which defines, for each gene *g*, a neighborhood *C*_*g*_ of genes with similar expression. The trended APL for gene *g* is the locally weighted neighborhood average,

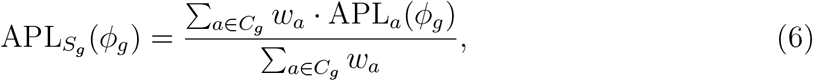

with tricube weights *w*_*a*_ (Cleveland, 1979) on AveLogCPM distance (Supplementary Note S6). The trended dispersion is 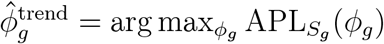.

##### Gene-specific (tagwise) dispersion

Each gene’s dispersion is shrunk toward the trended value via empirical Bayes (Smyth, 2004) by maximizing the weighted APL

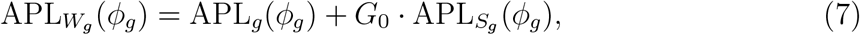

where *G*_0_ = *d*_0_*/d*_*g*_ is the empirical-Bayes prior weight estimated from the federated residual deviances at 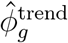. Larger *G*_0_ pulls 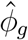 closer to 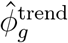. The gene-specific argmax is again located by natural cubic-spline interpolation through the 21 grid points. The full prior estimation and shrinkage procedure is shown in Supplementary Note S6.

#### 2.2.4 Likelihood ratio test

After estimating dispersions, we fit two models for each gene: a full model with all covariates and a reduced (null) model. Both fits use the federated IRLS procedure. Each client computes its local residual deviance 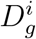 for both models. The deviance for the NB model is:

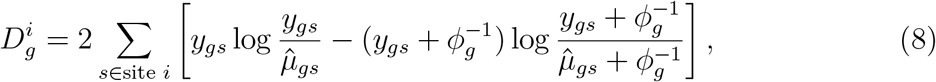

with the standard convention 0 log 0 ≡ 0 for the *y*_*gs*_ = 0 term, justified by 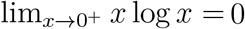. Since 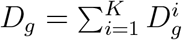, the total deviance is additive across sites and can be aggregated via SMPC. The aggregator then computes the likelihood ratio statistic, equivalently expressed via the maximized log-likelihoods or the deviance gap between models:

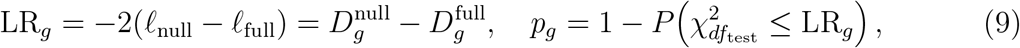

where ℓ_full_ and ℓ_null_ are the maximized NB log-likelihoods under the full and null designs. The term *df*_test_ is the difference in the ranks of the two design matrices. This reduces to the difference in column counts when both designs are full column-rank, the case in all our benchmarks. FDR-adjusted p-values are computed using the Benjamini-Hochberg procedure (Benjamini and Hochberg, 1995). See Supplementary Note S7 for the per-client log-likelihood decomposition and the privacy argument for the transmitted scalars.

### 2.3 Privacy analysis

FedEdgeR is a privacy-preserving framework built upon four security and architectural assumptions. First, all parties are honest-but-curious, meaning they adhere to the protocol but may attempt to infer private information from shared parameters. Second, the aggregator and compensator are non-colluding. Third, a minimum of three clients participate in the federation. Fourth, all components are deployed on physically distinct infrastructure. These assumptions align with existing federated frameworks such as Flimma (Zolotareva et al., 2021) and sPLINK (Nasirigerdeh et al., 2022), which are built on the HyFed architecture (Nasirigerdeh et al., 2021).

The SMPC masking protects data during transmission between parties. For integer parameters masked with uniform noise in ℤ_*p*_, the masked value 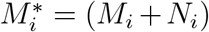 mod *p* is uniformly distributed regardless of *M*_*i*_, so the mutual information is zero: 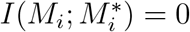 (Cramer et al., 2015). For real-valued parameters masked with Gaussian noise *N*_*i*_ ~ *N* (0, *σ*^2^), the information leakage is bounded by (Tjell and Wisniewski, 2021):

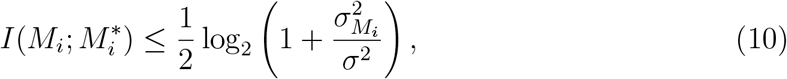

where 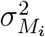 is the variance of the original parameter. With *σ*^2^ = 10^12^, this bound stays small for typical intermediate statistics (see Supplementary Notes S1 and S3 for the full derivation).

After denoising, the aggregator sees only global sums of per-gene per-site statistics; Supplementary Note S8 lists all transmitted quantities and their dimensions. When the number of samples per site is much larger than the number of covariates, recovering individual sample values from these aggregated statistics is not feasible.

The compensator sees only the random noise terms and never accesses any data or summary statistics. Raw count matrices and design matrices never leave the client sites. FedEdgeR does not provide formal differential privacy guarantees. While SMPC-based masking secures individual contributions during transmission, the aggregated statistics may still disclose sensitive information in scenarios with a very limited number of participants (e.g., *K* = 2). This limitation is shared by other federated tools like Flimma (Zolotareva et al., 2021) and sPLINK (Nasirigerdeh et al., 2022).

### 2.4 Compared methods

We compared FedEdgeR against four meta-analysis baselines. In each case, we ran edgeR independently at each client site and then combined the per-site results.

*Fisher’s method* (Fisher, 1934) *combines p-values using* 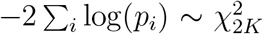. *Stouffer’s method* (Stouffer et al., 1949) converts p-values to z-scores (with sign from the log-fold change) and averages them. *RE model* uses the DerSimonian-Laird estimator (DerSimonian and Laird, 1986) to account for between-site heterogeneity in effect sizes. *RankProd* (Breitling et al., 2004) uses rank products to identify genes that are consistently ranked high across sites.

The gold standard for all comparisons is pooled edgeR. Pooled edgeR runs the standard edgeR pipeline on the aggregated data from all sites, as if the data were stored in a single repository.

### 2.5 Datasets

We evaluated FedEdgeR on four RNA-seq count datasets (Table 1) chosen to span four distinct evaluation regimes. BRCA provides a balanced binary contrast at moderate sample size. Mel-ICI is a multi-cohort contrast with study-of-origin covariates whose federation split follows natural cohort boundaries. GSE144269 is a binary tumor/non-tumor contrast on matched samples at moderate sample size in a non-TCGA cohort, fit with an unpaired two-group design (no per-patient effect). Oral is a high-dimensional small-sample stress test with an explicit paired design.

**Table 1:** Summary of the four benchmark datasets. BRCA and GSE144269 were partitioned into *K* = 3 client sites by deterministic class-stratified allocation. Each site preserves the global Normal-to-Tumor ratio. Oral was partitioned by positional slicing into three two-sample sites, with one patient’s matched tumor/normal pair on each site. Mel-ICI was partitioned along its natural cohort-of-origin boundaries (Gide / Hugo / Riaz), which correspond to three independent published studies.

| Dataset | #Genes | #Samples | Design cols. | Contrast |
| --- | --- | --- | --- | --- |
| BRCA | 14 864 | 227 | 2 | Tumor vs. Normal |
| Mel-ICI | 15 221 | 131 | 4 | Responder vs. non-responder (cohort-adjusted) |
| GSE144269 | 15 688 | 139 | 2 | Tumor vs. Normal |
| Oral | 10 514 | 6 | 4 | Tumor vs. Normal (paired, 3 patients) |

#### BRCA

A binary Tumor-vs-Normal subset of TCGA-BRCA (Weinstein et al., 2013) from the Genomic Data Commons. It was partitioned into *K* = 3 client sites by deterministic class-stratified allocation that preserves the per-site Normal-to-Tumor ratio. For the data-imbalance experiments in Section 3, two additional BRCA protocols are evaluated. The *size-imbalance* protocol allocates 70%*/*20%*/*10% of samples to the three sites while preserving the global Normal:Tumor ratio per site. The *ratio-imbalance* protocol keeps per-site sample counts roughly equal but skews the per-site Normal proportions to 72%*/*20%*/*4%.

#### Mel-ICI

A combined anti-PD-1 melanoma immune-checkpoint-inhibitor cohort assembled from three independent published RNA-seq studies (Gide et al., 2019, *n* = 56; Hugo et al., 2016, *n* = 26; Riaz et al., 2017, *n* = 49). The 4-column design (intercept, two cohort dummies, response indicator) tests response while adjusting for cohort of origin. Mel-ICI was partitioned along its natural cohort-of-origin boundaries (Gide / Hugo / Riaz), so each client corresponds to one of the three published studies.

#### GSE144269

The Mongolian hepatocellular carcinoma cohort published by Candia et al. (2020) (GSE144269) contains 70 tumor and 69 non-tumor liver samples, fit with an unpaired two-group design (no per-patient effect). It was partitioned into *K* = 3 client sites by deterministic class-stratified allocation that preserves the per-site Normal-to-Tumor ratio.

#### Oral

The three-patient paired oral squamous-cell carcinoma cohort of Tuch et al. (2010) is distributed with the NBPSeq R package (Di et al., 2011). With 6 samples and 4 design columns, the residual degrees of freedom are 2, making dispersion estimation sensitive to small numerical differences. Oral was partitioned by positional slicing into three two-sample sites, each holding one patient’s matched tumor/normal pair.

### 2.6 Implementation

FedEdgeR is implemented in Python 3.11 with R 4.5 invoked through rpy2 (≥3.6.7). The locfit-based (Loader, 1999) trended dispersion smoothing, the empirical-Bayes `squeezeVar` routine, and the `WLEB`-based per-gene tagwise argmax delegate to the original Bioconductor edgeR (≥4.8) implementation (Chen et al., 2025). The spline-refined per-gene tag-wise argmax is therefore identical to centralized edgeR’s default behavior. All federated arithmetic, IRLS iterations, and SMPC masking are pure Python (NumPy / SciPy). The three-party HyFed-style architecture (clients, aggregator, compensator) is realized here as process-isolated processes on a single host. A true distributed deployment with each role on a separate machine uses the same code paths and produces bit-for-bit identical results, because the SMPC masking and aggregation logic do not depend on the network transport. The benchmarks reported here were run on a single Linux server (AMD EPYC 7763, 64 cores, 528 GB RAM, RHEL 9.6). GPU acceleration is not used. We provide a reproducible Anaconda environment, deterministic random seeds for all stochastic steps, and an end-to-end driver script. Source code, data preparation scripts, and step-by-step reproducibility instructions are available at https://github.com/XunSong02/FedEdgeR.

## 3 Results

### 3.1 FedEdgeR reproduces pooled edgeR results

To evaluate performance, we first benchmarked FedEdgeR against the ground-truth pooled edgeR (executed on the aggregated dataset). Figure 2 presents the concordance of − log_10_(*p*-values) for all five methods on the four evaluation datasets.

**Figure 2:**
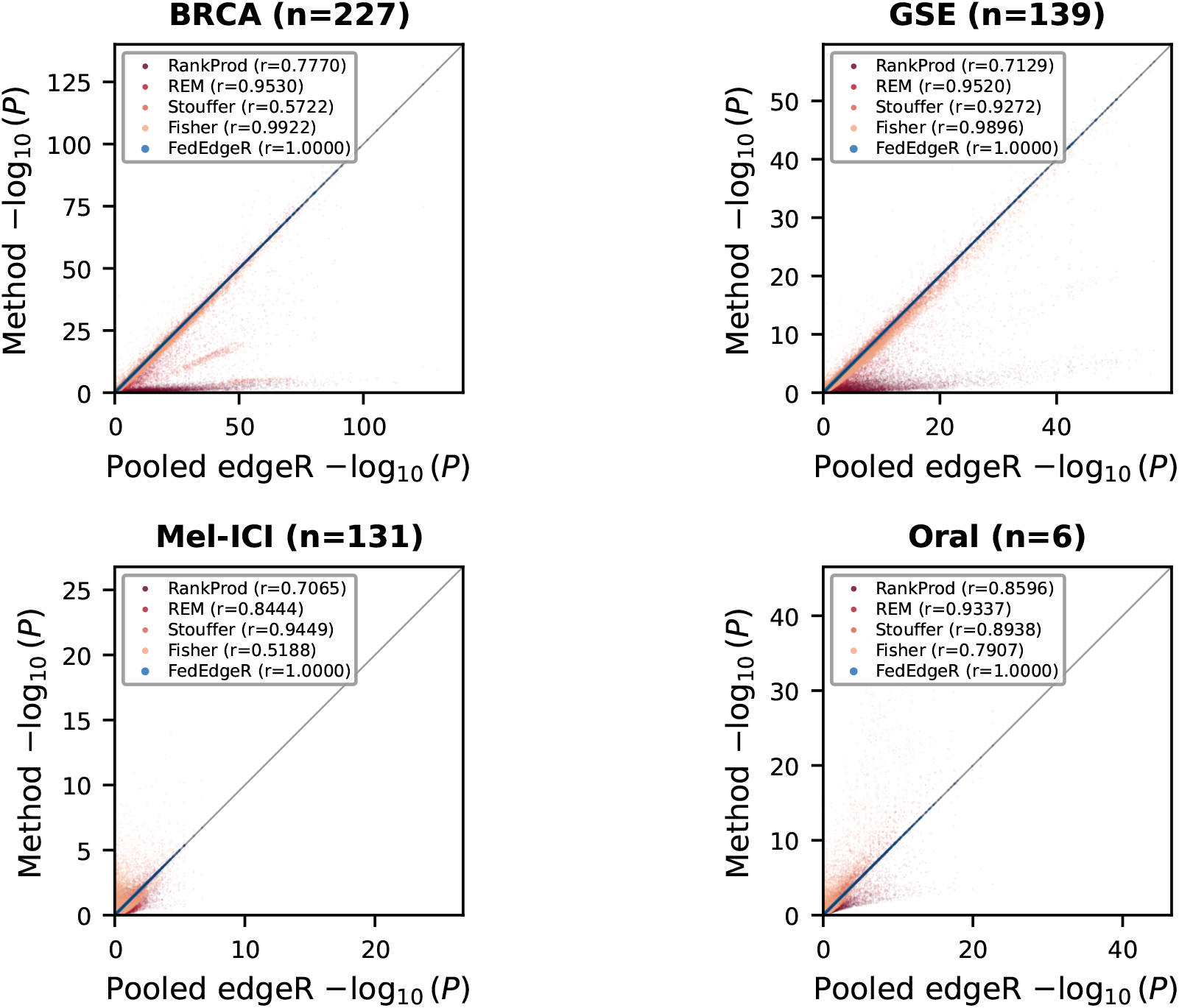
Concordance of −log_10_(*p*) between each method and pooled edgeR on four datasets. Each point is one gene. FedEdgeR (blue) falls on the diagonal (*y* = *x*), while meta-analysis methods scatter away from it. Pearson *r* is shown in each legend. **Alt text:** Four scatter plots in a 2 × 2 grid, one per benchmark dataset (BRCA *n* = 227, GSE *n* = 139, Mel-ICI *n* = 131, Oral *n* = 6). Each plot shows the −log_10_ *p*-value from one method versus pooled edgeR. FedEdgeR points (blue) lie on the *y* = *x* diagonal, indicating near-machine equivalence with pooled edgeR. Fisher, Stouffer, RE model, and RankProd points scatter away from the diagonal, with the magnitude of departure varying by dataset. Pearson *r* for each method is shown in the legend.

In the BRCA dataset (*n* = 227), FedEdgeR matched the pooled ground truth to near-machine precision. The Pearson correlation was 0.99999982 for log-fold changes (logFC) and 1.00000000 for *p*-values. The top 100 differentially expressed (DE) genes had 100% overlap with the pooled analysis. The F1 score for DE gene identification (FDR *<* 0.05) was 1.000.

The remaining three datasets showed the same pattern. On GSE144269 (*n* = 139), the *p*-value correlation, top 100 gene concordance, and F1 score were 1.00000000, 100%, and 1.000, respectively. The Mel-ICI dataset (*n* = 131) is qualitatively different. Each of the three federated clients corresponds to one of three independent published anti-PD-1 melanoma studies (Gide, Hugo, Riaz). The design carries study-of-origin dummy variables alongside the response indicator. Despite this naturalistic federation and the sparse DE signal (only 45 genes pass FDR*<* 0.05 in the pooled analysis out of 15,221 retained transcripts), FedEdgeR reproduced the centralized result. The *p*-value correlation reached 0.99999877, the top 100 overlap was 100%, and the F1 score was 1.000. The logFC correlation was 0.99906 (RMSE = 0.017). A small number of genes with very low residual expression in the cohort-adjusted contrast accounted for the largest per-gene logFC deviation (max |ΔlogFC| = 1.89). These genes were not in the significant set and did not affect either the ranking or the FDR call.

Oral (*n* = 6) has only two residual degrees of freedom under its four-column design. FedEdgeR still reached a *p*-value correlation of 1.00000000, 100% top-100 gene overlap, and an F1 score of 1.000. The logFC correlation was 0.99999 (RMSE = 0.005).

We further verified that the intermediate dispersion estimates generated by FedEdgeR are numerically consistent with pooled edgeR. EdgeR estimates dispersions at three levels. A single common dispersion is shared by all genes. A trended dispersion varies smoothly with average expression. A gene-specific (tagwise) dispersion is shrunk toward the trend via empirical Bayes (Smyth, 2004).

In the BRCA dataset, the common dispersion estimate differed by a negligible 1.4 *×* 10^−7^. Both trended and tagwise dispersions achieved Pearson correlations of 1.00000000, with RMSE values of 3.5 *×* 10^−8^ and 1.5 *×* 10^−7^, respectively.

On GSE144269, Mel-ICI, and Oral, high concordance was likewise maintained. Trended-dispersion Pearson correlations (*r*) were 1.00000000 in every case, while tagwise correlations ranged from 0.99999916 (Mel-ICI) to 1.00000000 (BRCA, GSE144269). The largest per-gene tagwise deviation, 1.6 *×* 10^−1^, occurred on a small subset of genes in Mel-ICI; on the remaining three datasets the maximum per-gene tagwise deviation stayed below 1.34 *×* 10^−4^. The Mel-ICI outlier is consistent with grid-induced discontinuities of the spline interpolant near nearly-flat APL maxima rather than a breakdown of the federated equivalence. The mechanism is detailed in Supplementary Note S6. The Mel-ICI common-dispersion drift is 1.3*×*10^−5^, versus ≤ 2.4*×*10^−7^ elsewhere. The shift is confined to a small subset of genes whose tagwise APL is flat near its maximum, none of which lie in the FDR-significant set. Top-100 overlap and F1 score on Mel-ICI both remain at 1.000.

The sufficient statistics decomposition (Eq. 4) ensures that FedEdgeR is theoretically equivalent to pooled edgeR under exact arithmetic. Although SMPC masking introduces floating-point rounding errors (typically near machine epsilon, ~ 10^−15^), these errors remain negligible even at the large masking variance *σ*^2^ = 10^12^. Our empirical results confirm that such numerical artifacts are inconsequential on all four benchmarks, including the smallest-sample Oral case.

### 3.2 FedEdgeR outperforms meta-analysis methods

Figure 2, Figure 3, and Table 2 present the performance comparison of FedEdgeR against the four classical meta-analysis methods.

**Table 2:** Quantitative comparison of FedEdgeR and meta-analysis methods against pooled edgeR. *r*: Pearson correlation. RMSE: root mean squared error of −log_10_(*p*). Top-100: overlap of the 100 most significant genes. F1: F1 score at FDR *<* 0.05. Bold indicates best per dataset.

| Dataset | Method | $r(\log\text{FC})$ | RMSE($-\log_{10} P$) | Top-100 | F1 |
| --- | --- | --- | --- | --- | --- |
| BRCA | FedEdgeR | **1.0000** | **0.000** | **1.00** | **1.000** |
|  | Fisher | 0.9982 | 1.867 | 0.85 | 0.952 |
|  | Stouffer | 0.9982 | 11.533 | 0.00 | 0.982 |
|  | RE model | 0.9974 | 4.366 | 0.69 | 0.978 |
|  | RankProd | 0.9982 | 16.289 | 0.23 | 0.127 |
| GSE | FedEdgeR | **1.0000** | **0.000** | **1.00** | **1.000** |
|  | Fisher | 0.9980 | 1.189 | 0.90 | 0.917 |
|  | Stouffer | 0.9980 | 2.466 | 0.26 | 0.981 |
|  | RE model | 0.9970 | 2.125 | 0.75 | 0.956 |
|  | RankProd | 0.9980 | 7.312 | 0.34 | 0.129 |
| Mel-ICI | FedEdgeR | **0.9991** | **0.0004** | **1.00** | **1.000** |
|  | Fisher | 0.8328 | 0.920 | 0.26 | 0.113 |
|  | Stouffer | 0.8328 | 0.211 | 0.64 | 0.481 |
|  | RE model | 0.8608 | 0.338 | 0.42 | 0.163 |
|  | RankProd | 0.8328 | 0.455 | 0.36 | 0.067 |
| Oral | FedEdgeR | **1.0000** | **0.000** | **1.00** | **1.000** |
|  | Fisher | 0.9950 | 2.646 | 0.32 | 0.709 |
|  | Stouffer | 0.9950 | 1.935 | 0.42 | 0.768 |
|  | RE model | 0.9961 | 0.657 | 0.59 | 0.878 |
|  | RankProd | 0.9950 | 1.324 | 0.26 | 0.324 |

**Figure 3:**
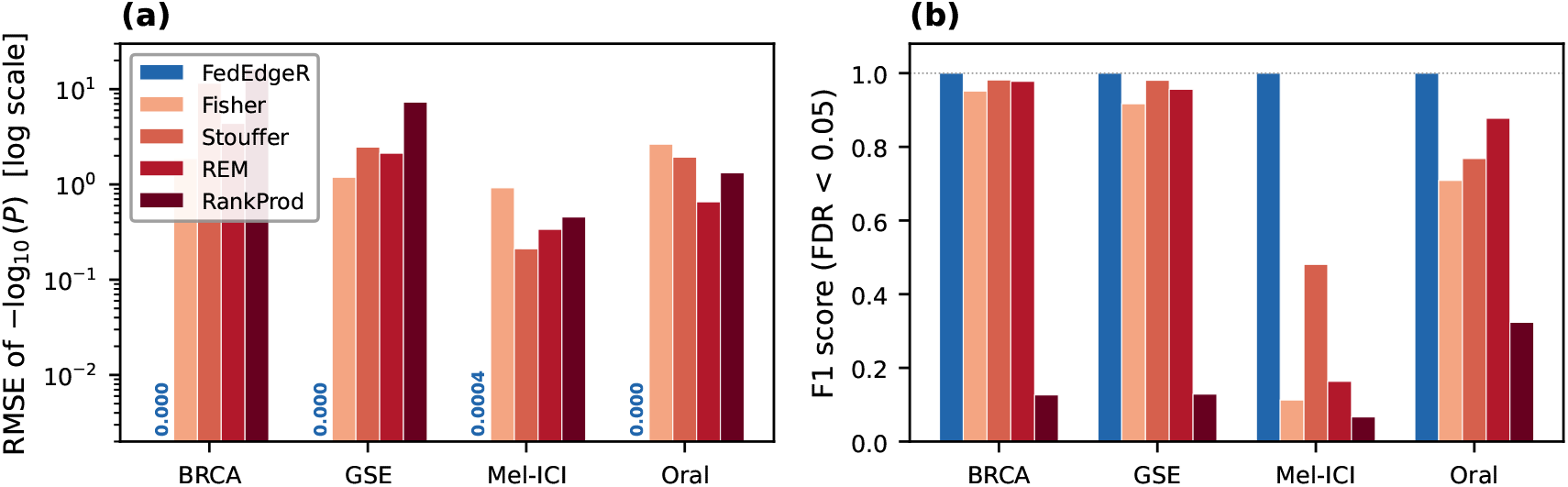
Comparison of FedEdgeR and four meta-analysis methods across datasets. (a) RMSE of −log_10_(*p*-values) relative to pooled edgeR (lower is better). (b) F1 score for DE gene identification at FDR *<* 0.05 (higher is better). **Alt text:** Two-panel bar chart comparing FedEdgeR with four meta-analysis baselines (Fisher, Stouffer, RE model, RankProd) across four datasets (BRCA, GSE, Mel-ICI, Oral). Panel (a): RMSE of −log_10_(*p*-values) on a log scale (lower is better). FedEdgeR’s bar is at zero on every dataset while baseline bars rise above. Panel (b): F1 score at FDR *<* 0.05 (higher is better). FedEdgeR reaches 1.000 on every dataset while baseline F1 varies, with RankProd consistently low.

FedEdgeR achieved the best performance on every evaluation metric. It had the lowest RMSE and the highest correlation and F1 scores on all four datasets. The F1 scores remained consistently at 1.000 across all benchmarking scenarios. Among meta-analysis baselines, Stouffer’s method was the strongest performer on the moderate-sample binary contrasts, with F1 of 0.982 on BRCA and 0.981 on GSE. However, Stouffer degraded sharply when the per-site signal was weak, with F1 dropping to 0.481 on Mel-ICI and 0.768 on Oral. The remaining baselines (Fisher, RE model, and RankProd) under-performed Stouffer on three of four datasets.

The performance disparity between FedEdgeR and the meta-analysis baselines was most pronounced on the Oral dataset. This gap is structural, not a consequence of tuning. After the *K* = 3 split, each site holds the matched tumor/normal pair of a single patient. The per-site design is 2 *×* 4 with rank 2 on 2 samples and zero residual degrees of freedom. Per-site edgeR therefore cannot estimate dispersion from the data, because the GLM is saturated at every gene. Producing any per-site *p*-value at all requires a rank reduction of the design plus a fixed-prior dispersion fallback at *ϕ* = 0.1, the edgeR initialization value. Supplementary Note S10 gives the per-site design matrices and the full fallback procedure. The Oral baseline numbers in Table 2 are therefore the output of a pipeline in which each site contributes information from a fixed-prior dispersion. FedEdgeR avoids this regime. The federated decomposition (Eq. 4) aggregates **X**^*T*^ **WX** and **X**^*T*^ **Wz** over all sites *before* any GLM is fit. Dispersion estimation therefore operates on the full six-sample design with df_res_ = 2 and matches pooled edgeR exactly. At *n* = 2 per site the meta-analysis route has no real per-site model left to fit, while sufficient-statistics aggregation reconstructs the pooled design without ever instantiating a per-site GLM.

RankProd underperformed on all four datasets (F1 0.067–0.324). Its non-parametric rank product is sensitive to noisy rankings on small per-site cohorts. Stouffer’s method achieved high F1 (≥ 0.981) on the strong-signal binary contrasts BRCA and GSE. However, its *p*-value-based rankings departed sharply from the pooled ground truth, with BRCA Top-100 overlap at 0% and GSE Top-100 at 26%. The combined z-scores inflate significance and create a large significant set with most true positives, but the within-set ranking does not track the pooled-edgeR signal. On Mel-ICI and Oral, Stouffer’s F1 dropped to 0.481 and 0.768, respectively.

### 3.3 Accuracy is stable across client counts

All previous experiments split each dataset across *K* = 3 client sites. To stress-test FedEdgeR’s numerical accuracy, we varied *K* from 2 to 10 (*K* = 2 is included only for numerical characterisation; privacy still requires *K* ≥ 3 per Section 2.3). Figure 4 shows that performance remained consistent across all tested *K*, so the *K* = 3 choice in earlier sections did not bias the result.

**Figure 4:**
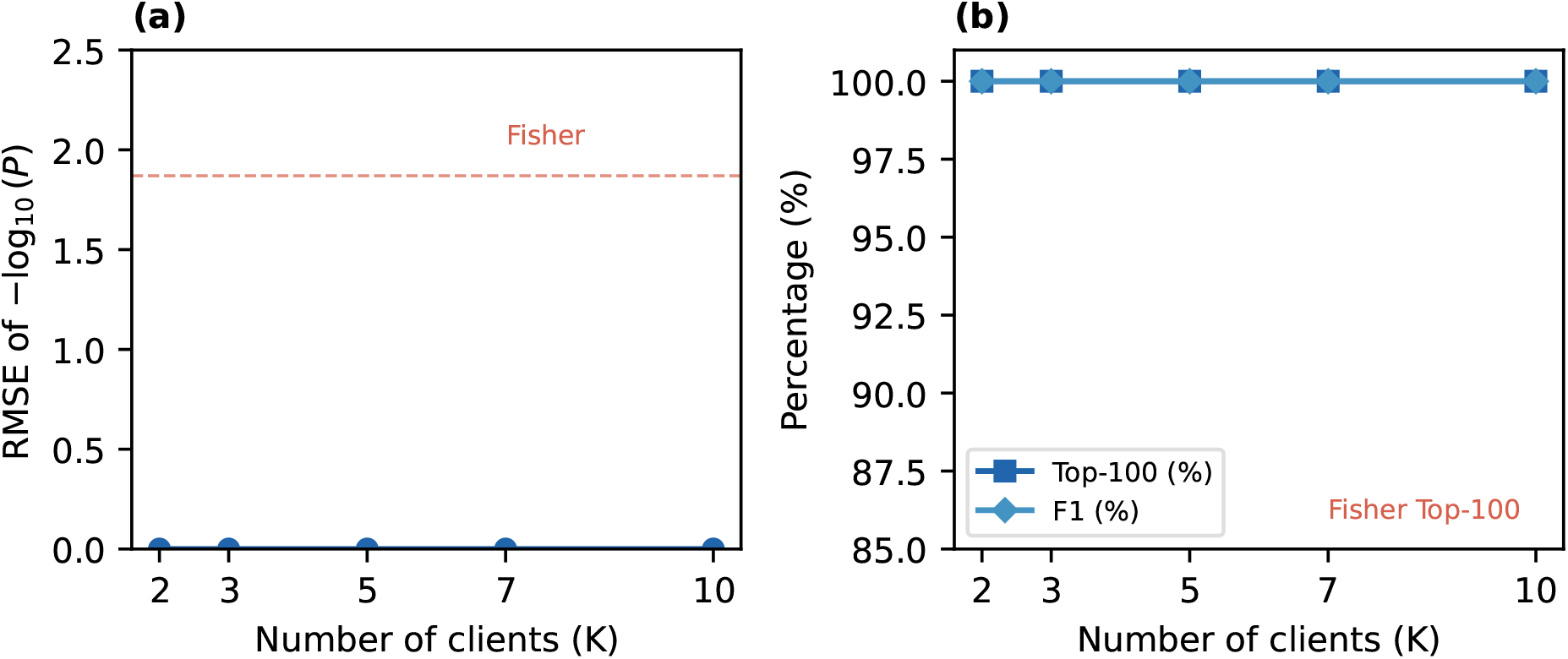
FedEdgeR accuracy on BRCA across *K* client sites. (a) RMSE of −log_10_(*p*). The red dashed line marks Fisher’s BRCA baseline (RMSE = 1.87), the best-performing meta-analysis baseline on this metric. (b) Top-100 gene overlap and F1 score. The red dashed line at 85% marks Fisher’s BRCA Top-100 overlap; F1 has no reference line because Stouffer (F1 = 0.982) outperforms Fisher (F1 = 0.952) on BRCA (cf. Table 2). **Alt text:** Two-panel line plot of FedEdgeR accuracy on BRCA as the number of client sites *K* varies from 2 to 10. Panel (a): RMSE of −log_10_(*p*) stays at zero for every *K*, well below Fisher’s BRCA baseline of 1.87 marked by a red dashed line. Panel (b): Top-100 gene overlap (squares) and F1 score (diamonds) remain at 100% and 1.000 across every *K*, well above Fisher’s 85% Top-100 reference (red dashed line). F1 has no reference line.

Over all tested values of *K*, FedEdgeR matched the centralized ground truth to near-machine precision. The Pearson correlation of logFC stayed at 0.99999983, while the top-100 DE gene overlap and F1 scores were 1.000 for every *K*. The RMSE of − log_10_(*p*) was also negligible (0.000) for all *K*. In contrast, Fisher’s method had a higher RMSE of 1.87 and a top-100 overlap of only 85%. Fisher is the most robust meta-analysis baseline on BRCA, yet it trails FedEdgeR.

We also tested imbalanced partitions on BRCA. The size-imbalance (70%*/*20%*/*10%) and ratio-imbalance (72%*/*20%*/*4% Normal proportions) protocols of Section 2 both yielded a *p*-value correlation of 1.000 and an F1 score of 1.000, identical to the balanced distribution. The decomposition (Eq. 4) makes the aggregated **X**^*T*^ **WX** and **X**^*T*^ **Wz** identical regardless of per-site allocation, so the local-to-global transition is lossless.

### 3.4 Runtime

End-to-end pipeline runtime ranges from approximately 21 seconds (Oral, *n* = 6) to 8.6 minutes (BRCA, *n* = 227) per dataset on a single workstation, dominated by the common-dispersion grid search. Per-step timings on the four benchmark datasets are reported in Supplementary Note S9.

## 4 Discussion

We presented FedEdgeR, a federated implementation of the edgeR pipeline for differential gene expression analysis. FedEdgeR uses SMPC to protect intermediate statistics during the multi-round IRLS fitting procedure. Our experiments on four real datasets showed that FedEdgeR produces results mathematically equivalent to a unified analysis. The p-value correlations and F1 scores were 1.000 on all four datasets. FedEdgeR therefore avoids the information loss of meta-analysis without sacrificing statistical power.

FedEdgeR addresses the remaining gap in federated differential expression (DE) analysis for bulk RNA-seq. While Flimma (Zolotareva et al., 2021) and FedPyDESeq2 (Muzellec et al., 2024) have federated the limma-voom and DESeq2 frameworks, respectively, edgeR remained the last major tool lacking a decentralized counterpart. These three methods are not functionally interchangeable. Whereas limma-voom models log-CPM values as continuous via linear regression, both edgeR and DESeq2 model count data directly using the negative binomial distribution. The federation of edgeR is inherently more complex than the single-pass weighted least squares (WLS) approach used in Flimma. It requires securing a multi-round iterative reweighted least squares (IRLS) procedure and the Cox-Reid adjusted profile likelihood (Cox and Reid, 1987). Because the corresponding routines are implemented in C inside the R edgeR package, the FedEdgeR client partially re-implements these subroutines in Python on top of HyFed. However, FedEdgeR preserves the statistical properties that make edgeR suitable for datasets with small sample sizes or high variability (Robinson and Smyth, 2007).

FedEdgeR achieved perfect concordance with the pooled ground truth even on the Oral dataset. Oral has only 6 samples, 4 covariates, and 2 residual degrees of freedom per gene. The top 100 differentially expressed genes were all recovered, and the F1 score was 1.000. The SMPC masking adds and subtracts Gaussian noise (*σ*^2^ = 10^12^) at each aggregation round. Although the noise cancels exactly in expectation, finite-precision arithmetic introduces rounding errors proportional to *σ* · *ε*_machine_ ≈ 10^−10^ per operation. These errors remain negligible on every dataset. The sufficient statistics decomposition therefore preserves full statistical equivalence in practice.

Communication rounds scale with the IRLS iteration count, since IRLS runs until the GLM coefficients converge. Per gene, each iteration transmits a *p × p* matrix and a *p ×* 1 vector, where *p* is the number of covariates and is typically 2 to 4 in our benchmarks. Because we batch all genes within each iteration, the per-round payload per client is approximately *G* · *p*^2^ · 8 bytes, where *G* is the number of retained genes. An additional *G* · *p* · 8 bytes carries the right-hand-side terms. For our four datasets the *p × p* block alone contributes roughly 0.5–2 MB per IRLS round per client. For example, *G*=15,221 and *p*=4 on Mel-ICI give ~ 1.95 MB for the *p × p* block. The *p ×* 1 vector adds at most another 0.5 MB, so the total per-round payload stays under 2.5 MB on every benchmark. Communication is not a bottleneck in practice, since each round completes in well under a second over standard networks.

FedEdgeR adopts the same honest-but-curious threat model as Flimma (Zolotareva et al., 2021) and sPLINK (Nasirigerdeh et al., 2022). The site-level decomposition handles arbitrary sample distributions without statistical penalty. The federated sufficient statistics depend only on the global **X**^*T*^ **WX** and **X**^*T*^ **Wz** pooled over all samples, regardless of how those samples are partitioned. Our size- and ratio-imbalance experiments confirm this property empirically.

The pipeline assumes that upstream pre-processing (read alignment, count generation, and normalization) is harmonized across sites. Heterogeneity at the count level is orthogonal to the federation problem and can be addressed with batch-effect correction tools such as ComBat (Leek et al., 2010) or federated batch correction methods such as FedscGen (Bakhtiari et al., 2025).

The pooled-equivalence numbers reported here use TMM normalization factors computed centrally on the merged per-site count matrices. Because pooled edgeR and FedEdgeR share the same upstream input, the equivalence claim isolates the federated DE pipeline. TMM’s M-value computation structurally requires the reference sample’s gene-level expression vector (Zolotareva et al., 2021). Federating TMM is therefore a separate open problem from federating edgeR’s DE pipeline. For end-to-end privacy in a real deployment, the upstream step can be replaced with Flimma’s federated UQ (Zolotareva et al., 2021). Flimma’s UQ exchanges only scalar normalization factors and never transmits raw counts. FedEdgeR accepts the resulting offsets unchanged, since its DE pipeline consumes per-sample offsets uniformly. We leave a fully federated TMM design to future work.

The most immediate next step is to integrate FedEdgeR into the FeatureCloud platform (Matschinske et al., 2023). Doing so would allow deployment in real multi-center studies without custom infrastructure. A natural follow-on is to add a differential privacy mechanism on top of the SMPC layer. Because IRLS iterates per gene, the privacy budget would need to compose over rounds, which trades off statistical accuracy against the formal privacy bound.

## Data availability

All datasets used in this study are publicly available. The BRCA dataset was derived from The Cancer Genome Atlas (TCGA-BRCA cohort) via the NCI Genomic Data Commons (GDC): https://portal.gdc.cancer.gov/projects/TCGA-BRCA. The Mel-ICI dataset combines three published anti-PD-1 melanoma RNA-seq studies, available under the following accessions: European Nucleotide Archive PRJEB23709 (https://www.ebi.ac.uk/ena/browser/view/PRJEB23709); NCBI Sequence Read Archive SRP067938 (https://www.ncbi.nlm.nih.gov/sra/?term=SRP067938) and SRP090294 (https://www.ncbi.nlm.nih.gov/sra/?term=SRP090294); and NCBI Sequence Read Archive SRP094781 (https://www.ncbi.nlm.nih.gov/sra/?term=SRP094781). The GSE144269 dataset is available from the NCBI Gene Expression Omnibus (GEO) under accession GSE144269: https://www.ncbi.nlm.nih.gov/geo/query/acc.cgi?acc=GSE144269. The Oral dataset is a three-patient paired oral squamous-cell carcinoma cohort, available as the oral carcinoma case-study data file accompanying the edgeR User’s Guide: https://bioinf.wehi.edu.au/edgeR/UserGuideData/TableS1.txt. All analysis code, conda environment, and reproducibility scripts are available at https://github.com/XunSong02/FedEdgeR. The version used in this manuscript is archived at Zenodo (https://doi.org/10.5281/zenodo.20450301).

## Author contributions

X.S. conceived the method, implemented the software, and wrote the manuscript. Z.W. supervised the research and revised the manuscript. Both authors read and approved the final manuscript.

## Acknowledgements

Anthropic’s Claude Opus 4.7 was used to assist with proofreading.

