## Supplementary for "FedEdgeR: federated and privacy-preserving edgeR for differential gene expression analysis"

**8 Supplementary Note S8. Per-step privacy summary 17**

**9 Supplementary Note S9. Runtime breakdown 18**

**10 Supplementary Note S10. Rank-deficient per-site designs and the small-cohort fallback for meta-analysis baselines 19**

**11 Supplementary Note S11. Use of large language models 21**

### 1 Supplementary Note S1. SMPC masking scheme

FedEdgeR uses the additive secret sharing scheme from HyFed (Nasirigerdeh et al., 2021), which traces back to the foundational secret sharing construction of Shamir (1979). We describe two variants: one for non-negative integer parameters and one for real-valued parameters.

#### 1.1 Non-negative integer parameters

The masking operates in the finite field $\mathbb{Z}_p = \{0, 1, \ldots, p-1\}$, where $p$ is a large prime (Cramer et al., 2015). We choose $p = 2^{54} - 33$, the largest prime that fits in a 54-bit integer.

Let $M_i$ denote the parameter held by client $i$, and let $N_i$ be a noise matrix drawn uniformly at random from $\mathbb{Z}_p$. The client computes the masked value:

$$M_i^* = (M_i + N_i) \bmod p.$$

The client sends $M_i^*$ to the aggregator and $N_i$ to the compensator.

The compensator computes the global noise:

$$N = \left( \sum_{i=1}^{K} N_i \right) \bmod p,$$

and sends $N$ to the aggregator.

The aggregator recovers the true global sum:

$$\begin{aligned} \left( \sum_{i=1}^{K} M_i^* - N \right) \bmod p &= \left( \sum_{i=1}^{K} (M_i + N_i) - \sum_{i=1}^{K} N_i \right) \bmod p \\ &= \left( \sum_{i=1}^{K} M_i \right) \bmod p. \end{aligned} \tag{1}$$

This is correct as long as $\sum_{i=1}^{K} M_i < p$. For typical genomic count data and library sizes, this condition is always satisfied with $p = 2^{54} - 33$.

In FedEdgeR's federated DE pipeline, the integer-mask variant is invoked only for the per-client maxima of the count matrix, $\max_{g,s} y^i_{gs}$. These maxima set the relative-tolerance scale for the IRLS convergence test in Algorithm S2. All other transmitted statistics listed in Table S1 are real-valued and use the Gaussian-mask variant of Section 1. The integer variant is retained as a primitive of the HyFed scheme. Future upstream extensions of FedEdgeR, such as a federated `filterByExpr`, can then reuse the same masking infrastructure without re-deriving privacy guarantees.

#### 1.2 Real-valued parameters

For real-valued parameters, the noise $N_i$ is drawn from a Gaussian distribution $\mathcal{N}(0, \sigma^2)$ with $\sigma^2 = 10^{12}$, following the same construction adopted by Flimma and sPLINK (Zolotareva et al., 2021; Nasirigerdeh et al., 2022). The masking is:

$$M_i^* = M_i + N_i.$$

The denoising follows the same additive cancellation:

$$\sum_{i=1}^{K} M_i^* - \sum_{i=1}^{K} N_i = \sum_{i=1}^{K} M_i.$$

Unlike the integer case, the denoised result is exact (no modular arithmetic), because the noise terms cancel perfectly.

#### 1.3 SMPC mask-and-denoise primitive

Algorithm S1 formalizes the mask-and-denoise procedure as a single primitive that all subsequent federated steps invoke. In the algorithm pseudocode that follows in this document, the line tag `[SMPC Aggregation]` stands for one invocation of Algorithm S1. The underlying noise generation, transmission to the compensator, noise-sum aggregation, and aggregator-side denoising are not repeated.

**Algorithm S1:** SMPC mask-and-denoise primitive (one aggregation round).

**Input:** Local value $M_i$ held by client $i \in \{1, \dots, K\}$.

**Data:** Noise model: $\mathcal{N}(0, \sigma^2)$ with $\sigma^2 = 10^{12}$ for real-valued $M_i$, or uniform $\mathbb{Z}_p$ with $p = 2^{54} - 33$ for non-negative integer $M_i$.

**Output:** Global sum $M = \sum_{i=1}^{K} M_i$ at the aggregator.

**for** *each client* $i = 1, \dots, K$ *in parallel* **do**

  sample noise $N_i$ from the chosen distribution;

  compute masked value $M_i^* \leftarrow M_i + N_i$;

  send $M_i^*$ to the aggregator;

  send $N_i$ to the compensator;

**end**

**compensator:** $N \leftarrow \sum_{i=1}^{K} N_i$, send $N$ to the aggregator;

**aggregator:** $M \leftarrow \left(\sum_{i=1}^{K} M_i^*\right) - N$;

**return** $M = \sum_{i=1}^{K} M_i$

### 2 Supplementary Note S2. Algorithm overview

Algorithm S3 states the FedEdgeR end-to-end pipeline. The pipeline follows the same conceptual stages as centralized edgeR: common dispersion via grid search, trended dispersion via locfit, tagwise dispersion via empirical Bayes, GLM-LRT, and BH-FDR. The federated IRLS sub-routine that fits a single negative-binomial GLM is factored out as Algorithm S2 and called by Algorithm S3 at every grid point of the common-dispersion search. It is also called once at $\hat{\phi}_g^{\text{trend}}$ to obtain the residual deviances consumed by `squeezeVar`, and for the full and null fits in Step 5. Per-step mathematical derivations are given in Supplementary Notes S4–S7; the per-step privacy reasoning is summarized in Supplementary Note S8.

In the pseudocode, line tags identify the actor performing each step:

- `[Client i]` denotes a per-site computation that runs in parallel at every client $i \in \{1, \dots, K\}$ on its own local data, with no data leaving the site.
- `[SMPC Aggregation]` denotes one invocation of Algorithm S1. The resulting global

sum is delivered at the aggregator.

- `[Aggregator]` denotes a server-side computation that uses only the SMPC-aggregated sums and, where applicable, broadcasts the result back to clients.

The compensator never appears in Algorithms S2 and S3. Its role is the noise-sum step inside Algorithm S1 and is absorbed into every `[SMPC Aggregation]` line.

**Algorithm S2:** Federated IRLS for one NB-GLM fit at fixed $\phi$ and design $\mathbf{X}$ (sub-routine).

**Input:** Per-client $(\mathbf{Y}^i, \mathbf{X}^i, \mathbf{o}^i)$ for $i = 1, \ldots, K$; dispersion $\phi$; tolerance $\varepsilon$; max iterations $T$.

**Output:** Coefficients $\hat{\boldsymbol{\beta}}_g$ for all genes $g$; per-site residual deviances $\{D_g^i\}_i$.

$t \leftarrow 0$; initialize $\boldsymbol{\beta}_g^{(0)}$ for all $g$;

**repeat**

  $t \leftarrow t + 1$;

  [Client $i$] compute $\mathbf{A}^i = (\mathbf{X}^i)^\top \mathbf{W}_g^i \mathbf{X}^i$;

  [Client $i$] compute $\mathbf{b}^i = (\mathbf{X}^i)^\top \mathbf{W}_g^i \mathbf{z}_g^i$;

  [SMPC Aggregation] $\mathbf{A} \leftarrow \sum_i \mathbf{A}^i$;

  [SMPC Aggregation] $\mathbf{b} \leftarrow \sum_i \mathbf{b}^i$;

  [Aggregator] $\boldsymbol{\beta}_g^{(t)} \leftarrow \mathbf{A}^{-1}\mathbf{b}$;

  [Aggregator] broadcast $\boldsymbol{\beta}_g^{(t)}$ to clients;

**until** $\max_g \|\boldsymbol{\beta}_g^{(t)} - \boldsymbol{\beta}_g^{(t-1)}\|_\infty < \varepsilon$ ***or*** $t = T$;

[Client $i$] compute residual deviance $D_g^i$ at $\hat{\boldsymbol{\beta}}_g \leftarrow \boldsymbol{\beta}_g^{(t)}$;

**return** $\hat{\boldsymbol{\beta}}_g$, $\{D_g^i\}_i$

**Algorithm S3:** FedEdgeR end-to-end pipeline. Step numbering here is finer-grained than in main-text Fig. 1. Steps 3 and 4 jointly correspond to its Tagwise stage, and Steps 5 and 6 to its GLM-LRT+BH-FDR stage.

**Input:** Per-client data $(\mathbf{Y}^i, \mathbf{X}^i, \mathbf{o}^i)$ for $i = 1, \ldots, K$; dispersion grid $\Phi = \{\phi_1, \ldots, \phi_{21}\}$; full and null design contrasts.

**Output:** Per-gene tagwise dispersion $\hat{\phi}_g^{\text{tag}}$, LR p-value $p_g$, BH-FDR $\text{FDR}_g$.

```
// Step 1: common dispersion (grid search over φ)
```

**foreach** $\phi_k \in \Phi$ **do**

> $\hat{\boldsymbol{\beta}}_g, \{D_g^i\}_i \leftarrow$ Algorithm S2 at $\phi_k$ under the full design;
>
> [Client $i$] compute $\ell_g^i(\phi_k; \hat{\boldsymbol{\beta}}_g)$;
>
> [Client $i$] compute $\mathcal{I}_g^i = (\mathbf{X}^i)^\top \mathbf{W}_g^i \mathbf{X}^i$;
>
> [SMPC Aggregation] $\ell_g \leftarrow \sum_i \ell_g^i$;
>
> [SMPC Aggregation] $\mathcal{I}_g \leftarrow \sum_i \mathcal{I}_g^i$;
>
> [Aggregator] $\text{APL}_g(\phi_k) \leftarrow \ell_g - \frac{1}{2} \log\det \mathcal{I}_g$;

**end**

[Aggregator] $\hat{\phi}_{\text{common}} \leftarrow \arg\max_\phi \sum_g \text{APL}_g(\phi)$;

```
// Step 2: trended dispersion
// Step 2a: federated Fisher-scoring for AveLogCPM (intercept-only
   NB-GLM)
```

initialize $\alpha_g$ for all $g$;

**repeat**

> [Client $i$] compute $U_g^i = \sum_{s \in \text{site } i} w_s (y_{gs} - \mu_{gs})/(1 + \mu_{gs}\phi_g)$;
>
> [Client $i$] compute $\mathcal{I}_g^i = \sum_{s \in \text{site } i} w_s \mu_{gs}/(1 + \mu_{gs}\phi_g)$;
>
> [SMPC Aggregation] $U_g \leftarrow \sum_i U_g^i, \quad \mathcal{I}_g \leftarrow \sum_i \mathcal{I}_g^i$;
>
> [Aggregator] $\alpha_g \leftarrow \alpha_g + U_g/\mathcal{I}_g$; broadcast to clients;

**until** $\alpha_g$ *converges for all* $g$;

[Aggregator] $\text{AveLogCPM}_g \leftarrow \log_2(e^{\alpha_g} \cdot 10^6)$;

```
// Step 2b: kernel-smooth APL across genes and per-gene argmax
```

[Aggregator] kernel-smooth $\text{APL}_g(\phi_k)$ across genes with tricube weights in AveLogCPM distance $\rightarrow \text{APL}_{S_g}(\phi_k)$;

[Aggregator] $\hat{\phi}_g^{\text{trend}} \leftarrow \arg\max_\phi \text{APL}_{S_g}(\phi)$;

```
// Step 3: refit at trended dispersion to obtain residual
```

### 3 Supplementary Note S3. Information leakage analysis

#### 3.1 Zero mutual information for integer parameters

For non-negative integer parameters masked with uniform noise in $\mathbb{Z}_p$, the masked value $M_i^* = (M_i + N_i) \bmod p$ is uniformly distributed over $\mathbb{Z}_p$ regardless of the value of $M_i$. This holds as long as $N_i$ is uniformly random and independent of $M_i$.

This means that $M_i$ and $M_i^*$ are statistically independent. The mutual information between the original and masked parameters is:

$$I(M_i; M_i^*) = 0.$$

An observer who sees only $M_i^*$ gains no information about $M_i$ (Cramer et al., 2015).

#### 3.2 Bounded mutual information for real-valued parameters

For real-valued parameters masked with Gaussian noise $N_i \sim \mathcal{N}(0, \sigma^2)$, the information leakage depends on the variance of the original parameter $M_i$ (Tjell and Wisniewski, 2021).

Assume $M_i$ has variance $\sigma_{M_i}^2$. The mutual information between $M_i$ and $M_i^* = M_i + N_i$ is maximized when $M_i$ is also Gaussian, by the maximum entropy property of the Gaussian distribution (Cover and Thomas, 2006). Under this worst-case assumption:

$$I(M_i; M_i^*) = \frac{1}{2} \log_2 \left(1 + \frac{\sigma_{M_i}^2}{\sigma^2}\right).$$

Since $M_i$ is generally not Gaussian, the actual mutual information satisfies:

$$I(M_i; M_i^*) \leq \frac{1}{2} \log_2 \left(1 + \frac{\sigma_{M_i}^2}{\sigma^2}\right).$$

With $\sigma^2 = 10^{12}$, the bound remains small even for statistics with large variance. For example, the IRLS working weights summed over 100 samples can produce $(\mathbf{X}^i)^T \mathbf{W}_g^i \mathbf{X}^i$ entries with $\sigma_{M_i}^2$ up to $\sim 10^8$, giving $I(M_i; M_i^*) \leq \frac{1}{2}\log_2(1+10^{-4}) \approx 7.2 \times 10^{-5}$ bits. This is negligible in practice.

### 4 Supplementary Note S4. Federated GLM fitting (glmFit)

#### 4.1 Overview

edgeR fits a negative binomial GLM (Nelder and Wedderburn, 1972) for each gene $g$ using iteratively reweighted least squares (IRLS) (Green, 1984; McCarthy et al., 2012; Chen et al., 2025). The model assumes $y_{gs} \sim \text{NB}(\mu_{gs}, \phi_g)$ with:

$$\text{E}(y_{gs}) = \mu_{gs}, \quad \text{Var}(y_{gs}) = \mu_{gs}(1 + \mu_{gs}\phi_g),$$

and the log-link function with per-sample offset $\log(\mu_{gs}) = \mathbf{x}_s^T \boldsymbol{\beta}_g + o_s$ (Robinson and Smyth, 2007), where $o_s = \log(N_s f_s)$ encodes the effective library size (raw library size $N_s$ times normalization factor $f_s$).

#### 4.2 IRLS iteration

At iteration $t$, given the current estimate $\boldsymbol{\beta}_g^{(t)}$, we compute for each sample $s$:

$$\mu_{gs}^{(t)} = \exp\left(\mathbf{x}_s^T \boldsymbol{\beta}_g^{(t)} + o_s\right), \tag{2}$$

$$w_{gs}^{(t)} = \frac{\mu_{gs}^{(t)}}{1 + \mu_{gs}^{(t)}\phi_g}, \tag{3}$$

$$z_{gs}^{(t)} = \mathbf{x}_s^T \boldsymbol{\beta}_g^{(t)} + \frac{y_{gs} - \mu_{gs}^{(t)}}{\mu_{gs}^{(t)}}. \tag{4}$$

The offset $o_s$ is included in $\mu_{gs}^{(t)}$ via the linear predictor, but does *not* appear in $z_{gs}^{(t)}$. This convention lets the closed-form weighted least-squares update below recover $\boldsymbol{\beta}_g^{(t+1)}$ directly without an offset subtraction.

The weight $w_{gs}$ comes from the variance function of the NB distribution. It equals $\mu_{gs}$ divided by the variance-to-mean ratio $1 + \mu_{gs}\phi_g$. The working variable $z_{gs}$ is the linearized version of the response (Green, 1984; McCarthy et al., 2012).

The update step is:

$$\boldsymbol{\beta}_g^{(t+1)} = \left(\mathbf{X}^T \mathbf{W}_g^{(t)} \mathbf{X}\right)^{-1} \mathbf{X}^T \mathbf{W}_g^{(t)} \mathbf{z}_g^{(t)},$$

where $\mathbf{W}_g^{(t)} = \mathrm{diag}(w_{g1}^{(t)}, \ldots, w_{gn}^{(t)})$.

#### 4.3 Federated decomposition

The matrix $\mathbf{X}^T\mathbf{W}_g\mathbf{X}$ is a $p \times p$ matrix, where $p$ is the number of covariates. We follow the additive decomposition strategy of Flimma and sPLINK (Zolotareva et al., 2021; Nasirigerdeh et al., 2022). The matrix can then be written as a sum over individual samples:

$$\mathbf{X}^T\mathbf{W}_g\mathbf{X} = \sum_{s=1}^{n} w_{gs}\, \mathbf{x}_s \mathbf{x}_s^T.$$

When samples are distributed across $K$ clients, this becomes:

$$\mathbf{X}^T\mathbf{W}_g\mathbf{X} = \sum_{i=1}^{K} \underbrace{\sum_{s \in \text{site } i} w_{gs}\, \mathbf{x}_s \mathbf{x}_s^T}_{(\mathbf{X}^i)^T \mathbf{W}_g^i \mathbf{X}^i}.$$

The same decomposition applies to $\mathbf{X}^T\mathbf{W}_g\mathbf{z}_g$:

$$\mathbf{X}^T\mathbf{W}_g\mathbf{z}_g = \sum_{i=1}^{K} (\mathbf{X}^i)^T \mathbf{W}_g^i \mathbf{z}_g^i.$$

Each client computes $(\mathbf{X}^i)^T\mathbf{W}_g^i\mathbf{X}^i$ (a $p \times p$ matrix) and $(\mathbf{X}^i)^T\mathbf{W}_g^i\mathbf{z}_g^i$ (a $p \times 1$ vector) locally. These are sent to the aggregator via SMPC masking. The aggregator sums them, inverts the $p \times p$ matrix, and obtains the new $\boldsymbol{\beta}_g$.

##### 4.4 Privacy of transmitted statistics

The aggregator receives $\mathbf{X}^T\mathbf{W}_g\mathbf{X}$ and $\mathbf{X}^T\mathbf{W}_g\mathbf{z}_g$. These are $p \times p$ and $p \times 1$ matrices aggregated across all $n$ samples.

In RNA-seq applications, $n$ is typically much larger than $p$. Sample counts can reach the hundreds, while $p$ is 2 to 4. Under this $n \gg p$ regime, the aggregated matrices have far fewer entries than the number of underlying sample-level values. Recovering the individual $w_{gs}$, $z_{gs}$, or $y_{gs}$ values from the aggregated matrices is a severely underdetermined problem (Zolotareva et al., 2021; Nasirigerdeh et al., 2022). The design matrix $\mathbf{X}$ is assumed to be known, since it encodes the experimental design rather than patient data. Even so, the system $\mathbf{X}^T\mathbf{W}\mathbf{z} = \mathbf{b}$ cannot be solved for the $n$ individual $(w_{gs}, z_{gs})$ pairs when $n \gg p$.

##### 4.5 Residual deviance

After convergence, the residual deviance for gene $g$ at site $i$ is (McCarthy et al., 2012):

$$D_g^i = 2 \sum_{s \in \text{site } i} \left[ y_{gs} \log \frac{y_{gs}}{\hat{\mu}_{gs}} - (y_{gs} + \phi_g^{-1}) \log \frac{y_{gs} + \phi_g^{-1}}{\hat{\mu}_{gs} + \phi_g^{-1}} \right].$$

The total deviance $D_g = \sum_{i=1}^{K} D_g^i$ is additive across sites.

#### 5 Supplementary Note S5. Federated adjusted profile likelihood

The adjusted profile likelihood (APL) for dispersion $\phi_g$, as employed by edgeR (McCarthy et al., 2012; Robinson and Smyth, 2007), is defined by

$$\text{APL}_g(\phi_g) = \ell(\phi_g; \mathbf{y}_g, \hat{\boldsymbol{\beta}}_g) - \frac{1}{2} \log \det(\mathcal{I}_g).$$

Here $\ell$ is the negative binomial log-likelihood and $\mathcal{I}_g = \mathbf{X}^T\mathbf{W}_g\mathbf{X}$ is the Fisher information matrix (the Cox-Reid adjustment term) (Cox and Reid, 1987).

#### 5.1 Log-likelihood decomposition

The NB log-likelihood for gene $g$ decomposes as a sum over samples (Robinson and Smyth, 2007):

$$\ell(\phi_g; \mathbf{y}_g, \hat{\boldsymbol{\beta}}_g) = \sum_{s=1}^{n} \ell_s(\phi_g; y_{gs}, \hat{\mu}_{gs}),$$

where each sample's contribution is:

$$\ell_s = y_{gs} \log \frac{\hat{\mu}_{gs}}{\hat{\mu}_{gs} + \phi_g^{-1}} + \phi_g^{-1} \log \frac{\phi_g^{-1}}{\hat{\mu}_{gs} + \phi_g^{-1}} + \log \binom{y_{gs} + \phi_g^{-1} - 1}{y_{gs}}.$$

When samples are distributed across $K$ sites:

$$\ell(\phi_g; \mathbf{y}_g, \hat{\boldsymbol{\beta}}_g) = \sum_{i=1}^{K} \ell^i(\phi_g; \mathbf{y}_g^i, \hat{\boldsymbol{\beta}}_g),$$

where $\ell^i$ is the sum of $\ell_s$ over samples at site $i$.

#### 5.2 Fisher information decomposition

The Fisher information matrix is:

$$\mathcal{I}_g = \mathbf{X}^T \mathbf{W}_g \mathbf{X} = \sum_{i=1}^{K} (\mathbf{X}^i)^T \mathbf{W}_g^i \mathbf{X}^i = \sum_{i=1}^{K} \mathcal{I}_g^i.$$

Both $\ell^i$ and $\mathcal{I}_g^i$ are computed locally by each client using the global $\hat{\boldsymbol{\beta}}_g$ and $\phi_g$ (broadcast by the aggregator) together with the local data $\mathbf{y}_g^i$ and $\mathbf{X}^i$. The local contributions are sent to the aggregator via SMPC masking, an additive aggregation strategy shared with other HyFed-based federated tools (Nasirigerdeh et al., 2021; Zolotareva et al., 2021; Nasirigerdeh et al., 2022). The aggregator then computes the full APL.

### 6 Supplementary Note S6. Federated dispersion estimation

#### 6.1 Common dispersion

The common dispersion $\phi$ is shared by all genes (Robinson and Smyth, 2007; McCarthy et al., 2012). It maximizes the total APL across genes,

$$\mathrm{APL}_C(\phi) = \sum_{g=1}^{G} \mathrm{APL}_g(\phi),$$

where $G$ is the total number of genes. The implementation sums the per-gene APL values. This is equivalent to maximizing the average, since the gene count $G$ is a positive constant that does not affect arg max.

FedEdgeR evaluates $\mathrm{APL}_C(\phi)$ at 21 grid points $\phi_k = 0.1 \cdot 2^{t_k}$, where $t_k$ are evenly spaced over $[-10, 10]$ in $\log_2$ space, so the grid spans $\phi \in [9.77 \times 10^{-5},\ 1.02 \times 10^{2}]$. At each grid point, the federated GLM fitting (Supplementary Note S4) is called to obtain $\hat{\boldsymbol{\beta}}_g$ for all genes. The federated APL (Supplementary Note S5) is then evaluated. The grid point that maximizes $\mathrm{APL}_C(\phi)$, identified by maximizing a natural cubic-spline interpolant through the 21 points, is selected as the common dispersion.

#### 6.2 Trended dispersion

The trended dispersion relates $\phi_g$ to the average expression level of each gene, measured by the average log-CPM (McCarthy et al., 2012; Chen et al., 2025). This step requires two federated computations.

*AveLogCPM computation.* Each gene's average log-CPM is computed through a federated Fisher-scoring iteration on the NB intercept-only mean model $\log \mu_{gs} = \alpha_g + o_s$. Here $o_s$ is the per-sample offset and $\phi_g$ is fixed to 0.05 to match edgeR's `aveLogCPM` default. At each iteration, every client sends the masked local score $U_g^i = \sum_{s \in \text{site } i} w_s (y_{gs} - \mu_{gs})/(1+\mu_{gs}\phi_g)$ and the Fisher information $\mathcal{I}_g^i = \sum_{s \in \text{site } i} w_s \mu_{gs}/(1+\mu_{gs}\phi_g)$ (with optional per-sample weights $w_s$ defaulting to 1). SMPC aggregation gives the global $U_g = \sum_i U_g^i$

and $\mathcal{I}_g = \sum_i \mathcal{I}_g^i$, and the aggregator updates $\alpha_g \leftarrow \alpha_g + U_g/\mathcal{I}_g$. Once $\alpha_g$ converges, the AveLogCPM is reported as $\log_2(e^{\alpha_g} \cdot 10^6)$ (the per-million log-rate, with library-size effects already absorbed into the offsets). We follow edgeR's default behavior (McCarthy et al., 2012; Chen et al., 2025). A small prior count (default 2, scaled by each sample's relative library size) is added to the counts and to the offsets before the Fisher-scoring iteration. This prevents log(0) instabilities for unexpressed genes. The shift depends only on per-sample library sizes and is computed locally by each client without exchanging counts. It does not require additional SMPC rounds.

*Local-regression fitting.* The aggregator obtains the gene-level APL values at each grid point and the AveLogCPM values. It then smooths the APL values across genes at every grid point, weighted by AveLogCPM distance. The implementation calls R's `locfit` (Loader, 1999) via the locfit option of edgeR's `WLEB` routine, which is edgeR's default trend smoother for `estimateDisp`. The trended dispersion for each gene is the argmax of its smoothed APL curve over the grid.

For each gene $g$, a neighborhood $C_g$ of genes with similar AveLogCPM values is selected. Tricube weights $w_a = (1 - |x_a|^3)^3$ (Cleveland, 1979) are then computed for each neighbor $a \in C_g$, where $x_a$ is the scaled difference in AveLogCPM. The weighted APL is:

$$\mathrm{APL}_{S_g}(\phi_g) = \frac{\sum_{a \in C_g} w_a \cdot \mathrm{APL}_a(\phi_g)}{\sum_{a \in C_g} w_a}.$$

#### 6.3 Gene-specific (tagwise) dispersion

Each gene receives its own dispersion by combining the gene-level APL with the trended prior. This follows the empirical Bayes shrinkage strategy of edgeR (Smyth, 2004; McCarthy et al., 2012; Robinson and Smyth, 2007).

First, the prior weight $G_0$ is estimated. Using the trended dispersion, the server computes the mean residual deviance $s_g^2 = D_g/d_g$ for each gene. These values are fitted to a scaled F-distribution $s_g^2 \sim s_0^2 \cdot F_{d_g,d_0}$ using the method of moments (Smyth, 2004). The prior degrees of freedom $d_0$ and prior mean $s_0^2$ are estimated from the empirical distribution of $s_g^2$. The prior weight is then:

$$G_0 = \frac{d_0}{d_g}.$$

The gene-specific dispersion maximizes:

$$\mathrm{APL}_{W_g}(\phi_g) = \mathrm{APL}_g(\phi_g) + G_0 \cdot \mathrm{APL}_{S_g}(\phi_g),$$

where $\mathrm{APL}_g$ is the gene-level APL and $\mathrm{APL}_{S_g}$ is the trended neighborhood APL. The term $G_0$ controls the strength of the shrinkage toward the trend. Larger $G_0$ (more prior information) pulls the gene-specific dispersion closer to the trended value (McCarthy et al., 2012). The argmax in $\mathrm{APL}_{W_g}(\phi_g)$ is located by a natural cubic-spline interpolant through the 21 $\log_2$ grid points. The result is $\hat{\phi}_g = 0.1 \cdot 2^{u_g^\star}$, where $u_g^\star$ is the spline argmax. The spline refinement is delegated to edgeR's `WLEB` routine via `rpy2` (Section 2.6 of the main text). The federated and centralized tagwise estimates therefore use the same per-gene continuous argmax.

**Grid-induced discontinuity on small APL maxima.** Although the spline refinement gives a continuous argmax in $u_g^\star$, the interpolant can have multiple near-equal local maxima when the underlying $\mathrm{APL}_{W_g}$ is flat near its peak. In that regime, a small floating-point increment can switch which local maximum dominates the natural cubic-spline argmax. This shifts $u_g^\star$ by roughly one grid step (step size 1.0 in $\log_2 \phi$) and produces a multiplicative $\sim 2$ factor in $\hat{\phi}_g$. This effect is amplified by larger baseline numerical drift in the federated IRLS path. On Mel-ICI, the 4-column cohort-adjusted design and the longer IRLS path produce a larger baseline drift than on the other three benchmark datasets.

### 7 Supplementary Note S7. Federated likelihood ratio test (glmLRT)

Once the final dispersion $\phi_g$ is obtained for each gene, the aggregator fits two models using the federated IRLS. The procedure follows the `glmLRT` test of edgeR (McCarthy et al., 2012; Chen et al., 2025):

- The **full model** includes all covariates in the design matrix $\mathbf{X}$ and produces fitted values $\hat{\mu}_{gs}^{\text{full}}$ and residual deviance $D_g^{\text{full}}$.
- The **null model** uses a reduced design matrix $\mathbf{X}_0$ (e.g., intercept only, or without the contrast of interest) and produces $\hat{\mu}_{gs}^{\text{null}}$ and $D_g^{\text{null}}$.

Each fit is performed by the federated IRLS routine of Supplementary Note S4, which already aggregates per-site residual deviances $D_g^{i,\text{full}}$ and $D_g^{i,\text{null}}$ via SMPC at the end of each fit:

$$D_g^{\text{full}} = \sum_{i=1}^{K} D_g^{i,\text{full}}, \quad D_g^{\text{null}} = \sum_{i=1}^{K} D_g^{i,\text{null}}.$$

The likelihood ratio statistic is the difference of total deviances. Recall that the residual deviance is defined as $D_g = -2(\ell_g - \ell_g^{\text{sat}})$, where $\ell_g^{\text{sat}}$ is the saturated log-likelihood evaluated at $\hat{\mu}_{gs} = y_{gs}$. This term depends only on the data, not on the fitted model. The saturated term therefore cancels in the difference:

$$\text{LR}_g = D_g^{\text{null}} - D_g^{\text{full}} = -2(\ell_{\text{null}} - \ell_{\text{full}}).$$

Under the null hypothesis, $\text{LR}_g$ approximately follows a $\chi^2$ distribution with $df_{\text{test}} = p_{\text{full}} - p_{\text{null}}$ degrees of freedom, where $p_{\text{full}}$ and $p_{\text{null}}$ are the number of parameters in the full and null models. The p-value is:

$$p_g = 1 - P\left(\chi^2_{df_{\text{test}}} \leq \text{LR}_g\right).$$

#### 7.1 Privacy of transmitted statistics

The aggregator receives $D_g^{\text{full}}$ and $D_g^{\text{null}}$, which are single scalar values per gene, each being a sum over all $n$ samples. These aggregated deviances reflect the overall model fit and do not contain enough information to reconstruct individual sample counts $y_{gs}$. The reasoning is the same as for the IRLS sufficient statistics in Supplementary Note S4 (Zolotareva et al., 2021; Nasirigerdeh et al., 2022).

### 8 Supplementary Note S8. Per-step privacy summary

Table S1 lists the statistics transmitted at each step of the FedEdgeR pipeline. It also gives their dimensions and the reason they do not leak individual sample data. Where applicable, we follow the privacy reasoning previously established for federated regression and GLM-based bioinformatics tools (Zolotareva et al., 2021; Nasirigerdeh et al., 2022, 2021).

Table S1: Summary of transmitted statistics in the federated phase of FedEdgeR. Filtering and TMM normalization sit upstream of the federated pipeline. In this benchmark the TMM factors were computed centrally on the pooled per-site count matrices (Methods §2.2.1, Discussion).

| Step | Transmitted statistic | Dimension | Why safe |
|---|---|---|---|
| glmFit (per iteration) | $(\mathbf{X}^i)^T\mathbf{W}_g^i\mathbf{X}^i$ | $p \times p$ | Aggregated over $n_i$ samples with $n_i \gg p$ |
| | $(\mathbf{X}^i)^T\mathbf{W}_g^i\mathbf{z}_g^i$ | $p \times 1$ | Same argument |
| APL | $\ell^i(\phi_g; \mathbf{y}_g^i, \hat{\boldsymbol{\beta}}_g)$ | scalar | Sum over $n_i$ samples |
| | $\mathcal{I}_g^i$ | $p \times p$ | Same as glmFit |
| AveLogCPM | Score $U_g^i$, Fisher info $\mathcal{I}_g^i$ | scalar each | Sums over samples |
| LRT | $D_g^{i,\text{full}}, D_g^{i,\text{null}}$ | scalar each | Sums over samples (sent at the end of each federated IRLS fit) |
| Tagwise refit | $D_g^i$ at $\hat{\phi}_g^{\text{trend}}$ | scalar | Sum over samples, consumed by squeezeVar |

In all cases, the transmitted quantities are sums or weighted sums over all samples at a client site. The number of samples per site ($n_i$) is much larger than the number of model parameters ($p$). Under this condition, reconstructing individual sample values from these aggregated statistics is an underdetermined problem (Zolotareva et al., 2021; Nasirigerdeh et al., 2022). The SMPC masking provides an additional layer of protection during transmission.

### 9 Supplementary Note S9. Runtime breakdown

We measured the wall-clock cost of each pipeline step on each of the four benchmark datasets. The benchmarks ran on a single Linux server (AMD EPYC 7763, 64 cores, 528 GB RAM, RHEL 9.6) in the single-host multi-process simulation mode (see Implementation in the main text). Common-dispersion estimation dominates the total cost because it evaluates the federated APL at 21 dispersion grid points, and each grid-point evaluation invokes a full IRLS fit. The trended-dispersion, tagwise-dispersion, GLM-LRT, and top-tags export steps together account for between 4 s (Oral) and 89 s (BRCA). This is at most about 19% of end-to-end runtime in any dataset.

Table S2: Per-step wall-clock time (seconds) for FedEdgeR with $K = 3$ on the four benchmark datasets, measured during the unified end-to-end runs reported in the main text. Sample sizes are BRCA $n = 227$, GSE144269 $n = 139$, Mel-ICI $n = 131$, and Oral $n = 6$. The total time is the sum of all listed steps. The GLM-LRT step covers the final full-vs-null IRLS fit and the likelihood-ratio test.

| Step | BRCA | GSE144269 | Mel-ICI | Oral |
|---|---|---|---|---|
| Common dispersion (grid×IRLS) | 428.7 | 244.0 | 457.7 | 16.8 |
| Trended dispersion (locfit) | 1.9 | 1.3 | 2.8 | 0.6 |
| Tagwise dispersion (EB shrinkage) | 34.1 | 15.0 | 15.7 | 0.9 |
| GLM fit + LRT | 52.4 | 28.2 | 37.5 | 2.0 |
| Top-tags export | 0.2 | 0.2 | 0.4 | 0.2 |
| End-to-end (per run) | 517.7 | 289.0 | 514.8 | 20.8 |

Three observations about the timing pattern. First, common-dispersion estimation dominates total runtime in every dataset (81–89% of end-to-end), because the 21-point grid search invokes a full IRLS fit at each grid point. Second, runtime scales jointly with sample count $n$ and design-matrix width $p$. BRCA (large $n$, small $p$) and Mel-ICI (moderate $n$, $p$=4) take comparable total time, while GSE144269 (moderate $n$, small $p$) is roughly half as expensive. Oral ($n = 6$) finishes in about 21 s because its tiny sample count keeps every per-IRLS matrix small. Third, in all four datasets the per-round SMPC communication payload is at most a few hundred bytes per gene (a $p \times p$ matrix and a $p \times 1$ vector with $p \in \{2, 3, 4\}$). Network transport is therefore not a bottleneck. Under realistic multi-site network conditions the wall-clock time is dominated by computation rather than communication.

### 10 Supplementary Note S10. Rank-deficient per-site designs and the small-cohort fallback for meta-analysis baselines

The Oral cohort (Tuch et al., 2010) contains six samples drawn from three patients, with each patient contributing one matched tumor/normal pair. The global design used in the main-text analyses has four columns (intercept, two patient indicators, and one tumor indicator). The residual degrees of freedom are therefore $\mathrm{df}_{\mathrm{res}}^{\mathrm{global}} = 6 - 4 = 2$. After the $K = 3$ federated split, each of the three sites receives a single patient's matched pair. The per-site design matrices on the two-sample sub-cohorts are

$$\mathbf{X}^{(1)} = \begin{pmatrix} 1 & 0 & 0 & 0 \\ 1 & 0 & 0 & 1 \end{pmatrix}, \quad \mathbf{X}^{(2)} = \begin{pmatrix} 1 & 1 & 0 & 0 \\ 1 & 1 & 0 & 1 \end{pmatrix}, \quad \mathbf{X}^{(3)} = \begin{pmatrix} 1 & 0 & 1 & 0 \\ 1 & 0 & 1 & 1 \end{pmatrix},$$

where the columns are ordered (intercept, patient-2 indicator, patient-3 indicator, tumor indicator). Each $\mathbf{X}^{(i)}$ is $2 \times 4$ with $\mathrm{rank}\big(\mathbf{X}^{(i)}\big) = 2$. On every site one patient column is identically zero and the other is collinear with the intercept. The relevant patient is either absent from the site or present in both rows. The per-site residual degrees of freedom are therefore

$$\mathrm{df}_{\mathrm{res}}^{(i)} \; = \; n_i - \mathrm{rank}\big(\mathbf{X}^{(i)}\big) \; = \; 2 - 2 \; = \; 0.$$

**Consequences for meta-analysis baselines.** At $\mathrm{df}_{\mathrm{res}}^{(i)} = 0$ the per-site GLM is saturated. The negative-binomial likelihood is maximised by the data themselves. The residual deviance therefore vanishes for every gene $g$, and the dispersion $\phi$ is not identifiable from the per-site fit. Two adjustments are unavoidable before any per-site $p$-value can be produced and passed to a meta-analysis combiner.

1. *Rank reduction of the per-site design.* A QR factorization with column pivoting selects $\mathrm{rank}\big(\mathbf{X}^{(i)}\big) = 2$ linearly independent columns and drops the remaining two. The contrast vector $\mathbf{c} = (0, 0, 0, 1)^T$ is reduced consistently with the pivot. After reduction, the per-site design retains the intercept and the tumor indicator on every site. The tumor indicator is the only contrast each site can estimate.

2. *Constant-dispersion fallback.* With $n_i \leq p_i$ even after rank reduction, the Cox-Reid grid search and the empirical-Bayes shrinkage that would normally drive `estimateDisp` have no residual deviance to consume. We therefore set $\phi_g^{(i)} = 0.1$ for all genes and all sites, the value used by `edgeR::DGEList` as its initial common-dispersion estimate. We then proceed with `glmFit` and `glmLRT`. The constant 0.1 is a fixed prior. It is not estimated from the per-site data. The baseline driver `run_baselines.R` records the fallback explicitly in its per-cohort log.

The per-site $p$-values produced under these adjustments are then combined by Fisher's method, Stouffer's method, the RE model, and RankProd. The combiners are exactly those described in Section 2.4 (Compared methods) of the main text. The Oral baseline numbers reported in Table 2 of the main text (Fisher F1 = 0.709, Stouffer F1 = 0.768, RE model F1 = 0.878, RankProd F1 = 0.324) are the output of this rank-reduced, fixed-dispersion pipeline. Without the two adjustments above, the per-site edgeR fit aborts at the dispersion-estimation step and no $p$-value is emitted. Meta-analysis on three $n = 2$ sites would then have no input to combine at all.

**Why FedEdgeR is unaffected.** FedEdgeR never instantiates a per-site GLM. The sufficient-statistics decomposition (Eq. 4 of the main text) sums the per-site weighted normal equations

$$\mathbf{X}^T\mathbf{W}\mathbf{X} \;=\; \sum_{i=1}^{K}\mathbf{X}^{(i)\,T}\mathbf{W}^{(i)}\mathbf{X}^{(i)}, \qquad \mathbf{X}^T\mathbf{W}\mathbf{z} \;=\; \sum_{i=1}^{K}\mathbf{X}^{(i)\,T}\mathbf{W}^{(i)}\mathbf{z}^{(i)},$$

*before* any IRLS solve is performed. The aggregated $\mathbf{X}^T\mathbf{W}\mathbf{X}$ has rank 4, the same rank as the global Oral design. The federated IRLS, the federated APL grid search, and the federated tagwise empirical-Bayes step therefore all operate on the full six-sample design with $\mathrm{df}_{\mathrm{res}} = 2$. The per-site rank deficiency observed in $\mathbf{X}^{(i)}$ above is invisible to the federated pipeline. Dispersion is estimated from a pooled adjusted profile likelihood. Per-site residual deviances are never instantiated.

**Take-away.** The Oral comparison illustrates a structural rather than empirical advantage of sufficient-statistics federation over meta-analysis. As the per-site cohort size shrinks, meta-analysis approaches a regime in which the per-site model is unidentifiable. Constant-dispersion fallbacks make the pipeline runnable but propagate information loss that no downstream $p$-combiner can repair. Sufficient-statistics aggregation does not encounter this regime, because the model that is fit is, by construction, the pooled model on $\sum_i n_i$ samples. The Oral row of Table 2 should therefore be read as a demonstration of this structural difference. At $n = 2$ per site, meta-analysis baselines are running on a degraded input that they cannot avoid. FedEdgeR is running on the same data the centralized pipeline would have seen.

### 11 Supplementary Note S11. Use of large language models

This manuscript and its supplementary materials follow the International Society for Computational Biology (ISCB) acceptable use policy for large language models (LLMs) endorsed by *Bioinformatics*. The authors used Anthropic's Claude Opus 4.7 as a writing aid for grammar and clarity polishing of text that the authors had drafted. The model edited individual sentences and reorganized paragraphs at the authors' direction. Every edit was reviewed and accepted, modified, or rejected by the authors.

The model was not used to draft any section from a prompt, to generate the manuscript figures, to write any portion of the FedEdgeR source code, to generate or analyse data, or to formulate scientific claims, derivations, experimental results, or their interpretation. The model is not listed as an author and does not satisfy ICMJE authorship criteria.
